# tDCS Cranial Nerve Co-Stimulation: Unveiling Brainstem Pathways Involved in Trigeminal Nerve Direct Current Stimulation in Rats

**DOI:** 10.1101/2024.10.11.617809

**Authors:** Alireza Majdi, Lars E. Larsen, Robrecht Raedt, Myles Mc Laughlin

## Abstract

The effects of transcranial direct current stimulation (tDCS) are typically attributed to the polarization of cortical neurons by the weak electric fields it generates in the cortex. However, emerging evidence indicates that certain tDCS effects may be mediated through the co-stimulation of peripheral or cranial nerves, particularly the trigeminal nerve (TN), which projects to critical brainstem nuclei that regulate the release of various neurotransmitters throughout the central nervous system. Despite this, the specific pathways involved remain inadequately characterized.

In this study, we examined the effects of acute transcutaneous TN direct current stimulation (TN-DCS) on tonic (i.e. mean spike rate and spike rate over time) and phasic (number of bursts, spike rate per burst, burst duration, and inter-burst interval) activities while simultaneously recording single-neuron activity across three brainstem nuclei in rats: the locus coeruleus (LC), dorsal raphe nucleus (DRN), and median raphe nucleus (MnRN).

We found that TN-DCS significantly modulated tonic activity in the LC, with notable interactions between stimulation amplitude, polarity, and time epoch affecting mean spike rates. Similar effects were observed in the DRN regarding tonic activity. Further, phasic activity in the LC was influenced by TN-DCS, with changes in burst number, duration, and inter-burst intervals linked to stimulation parameters. Conversely, MnRN tonic activity following TN-DCS remained unchanged. Importantly, xylocaine administration to block TN abolished the effects on tonic activities in both the LC and DRN.

These results suggest that tDCS effects may partially arise from indirect modulation of the TN, leading to altered neuronal activity in DRN and LC. Besides, the differential changes in tonic and phasic LC activities underscore their complementary roles in mediating TN-DCS effects on higher cortical regions. This research bears significant translational implications, providing mechanistic insights that could enhance the efficacy of tDCS applications and deepen our understanding of its neurophysiological effects.

## Introduction

Alterations in central noradrenergic (NA) and serotonergic system functioning are critical in the onset and progression of neuropsychiatric disorders (1). The majority of central NA neurons are located in the locus coeruleus (LC)(2), whereas the dorsal and median raphe nuclei (DRN and MnRN, respectively) house the highest concentrations of serotonergic neurons in the brain. These neurotransmitter systems have been proposed as potential neuromodulation targets for boosting cognitive function (3, 4) and treating neuropsychiatric disorders (5, 6). However, the deep location of these nuclei in the brainstem poses challenges for direct stimulation using non-invasive neuromodulation methods.

Transcranial direct current stimulation (tDCS) is a noninvasive neuromodulation method that uses scalp electrodes to deliver direct currents of 1 to 5 mA, creating a weak electric field in the brain (7). It is typically assumed that the neuromodulatory effects of tDCS are solely due to this weak electric field generated in the cortex. However, recent evidence suggests that some tDCS effects may be attributed to the co-stimulation of nerves in the scalp, such as the trigeminal nerve (TN) and the occipital nerve (8, 9). Following TN stimulation, the trigeminal principal sensory nucleus (NVsnpr) and the mesencephalic nucleus (MeV) are activated (10). These trigeminal nuclei have extensive connections with the serotonergic and NA nuclei, thereby modulating neural activity in higher cortical areas such as the medial prefrontal cortex (11) and the hippocampus (12–14). Despite this potential, the effects of TN direct current stimulation (TN-DCS) on these pathways have not been previously studied.

LC exhibits two distinct firing modes: phasic and tonic. Phasic LC activity, characterized by brief bursts of high-frequency firing, is often triggered by salient stimuli and is thought to enhance attention and facilitate responses to such stimuli (15). In contrast, tonic LC activity, a more sustained, baseline firing level, is associated with general arousal and vigilance (16). This tonic mode modulates the brain’s overall responsiveness and preparedness to react (17). Yet, these two modes of activity can dynamically interact to optimize behavioral performance in response to environmental demands (18). In line with that, LC-NA neurons display an increased firing rate followed by a subsequent period of suppressed activity in response to pulsed peripheral nerve stimulation (19). Notably, the LC is recognized as a crucial mediator of vagus nerve stimulation (VNS) effects within the central nervous system (20, 21). Previous studies have demonstrated that VNS with pulse trains lasting a few seconds induces an elevation in firing rates and an increase in the percentage of neurons firing in bursts in the LC. This response develops over minutes to hours (22, 23). However, the effects of TN-DCS on any of these firing modes are not yet known.

The DRN and MnRN comprise 5-hydroxytryptaminergic (5HT) and non-5HT neurons (24–26). The 5-HT neurons within the DRN demonstrate specific projection patterns, with the caudal DRN primarily projecting to the hippocampus (27). Evidence has revealed that chronic pulsed VNS substantially increases the firing rate of 5HT neurons in the DRN compared to control rats (23). Furthermore, the MnRN serves as the primary source of extensive 5-HT innervation to complete hippocampal formation (24). Evidence suggests that distinct neuronal connections, i.e. serotonergic and glutamatergic, within the MnRN-hippocampal pathway, facilitate different brain states (e.g. non-theta and theta, respectively) (28). This implies that a non-serotonergic system originating from the MnRN, possibly glutamatergic, may play a distinct role apart from the ascending serotonergic raphe projection in regulating hippocampal network activity (28).

In prior studies (10, 14), our research group was the first to demonstrate a significant increase in neural activity in the trigeminal nuclei, i.e., the NVsnpr and MeV, in rats following TN-DCS. Building upon this foundation, the present study aimed to further elucidate the mechanism of the action of TN-DCS by investigating its effect on neuronal activity in the LC, DRN, and MnRN.

## Materials and methods

### Subjects, anesthesia, and surgery

A total of 21 adult male Sprague Dawley rats obtained from Charles River Laboratories were utilized for this study. The allocation comprised 9 rats for the RNs group and 12 for the LC group (7 rats cathodic vs. 11 rats for anodic stimulation). Before surgical procedures, rats weighing 250-350 g were carefully paired and housed in cages under tightly controlled environmental conditions, maintaining a diurnal light-dark cycle. During the pre-experimental phase, rats were provided unrestricted access to food and water. It is essential to emphasize that experiments were terminal. Adherence to the ARRIVE guidelines and compliance with the EU Directive 2010/63/EU governing animal experiments were consistently ensured throughout all animal-related procedures. Experimental protocols were approved by the KU Leuven Ethics Committee for laboratory experimentation under license numbers 201/2018 and 072/2020.

On experiment days, rats were anesthetized via intraperitoneal (i.p.) injection of urethane (ethyl carbamate; Sigma-Aldrich, U2500) at a dose of 1.5 g/kg body weight (2) and positioned in a stereotaxic frame (RWD 71000 Automated Stereotaxic Instrument, RWD Life Science Co., Ltd, Shenzhen, China) on a heating pad. The core body temperature of the rats was monitored and maintained at approximately 37 ± 0.5 °C using a plastic rectal probe (ThermoStar Homeothermic Monitoring System, RWD Life Science Co., Ltd, Shenzhen, China). Anesthesia depth was regularly evaluated by observing the toe-pinch reflex, with adjustments made through additional urethane injection (approximately 100 µL via i.p. route) to maintain a consistent level (2). Following scalp removal, meticulous skull adjustments were made to achieve a level of 0 degrees on both the anteroposterior and mediolateral planes, ensuring less than 0.2mm differences between lambda-bregma and lateral ridges.

### Stereotaxic coordinates and electrode placement

Figure 1 illustrates the sagittal plane of the rat brain, depicting the relative positions of the recording and stimulating electrodes, as well as the TN and its primary nuclei, the LC, DRN, and MnRN.

Neural activity in the LC was monitored using a 32-channel silicon probe (Cambridge Neurotech, Cambridge, UK, Model: ASSY-37-H6B). The craniotomy was performed with the 78001 Microdrill (RWD Life Science Co., Shenzhen, China). The LC electrode, implanted at a 15-degree posterior angle, was positioned at stereotaxic coordinates 3.9-4.2 mm posterior and 0.6-1.2 mm lateral to true lambda, and approximately 6.0 to 6.5 mm ventral from the brain surface. The placement of the LC electrode adhered to established electrophysiological and anatomical criteria (2).

Neural activity was carefully monitored across all 32 channels during electrode insertion. The array was gradually advanced ventrally until the characteristic biphasic response of LC-NA neurons—marked by excitation followed by auto- and lateral inhibition—was triggered by a noxious foot shock (3.0 - 4.0 mA biphasic square pulse, 0.1 msec duration) (29). Neurons having >2% of recorded spikes within a predetermined 3 ms refractory period were removed to guarantee that single units were being evaluated (30–33). During the designated period, spike waveforms were extracted from the electrode contact displaying the highest amplitude of the neuron. The characteristics of LC neurons were assessed using three primary metrics: mean spike rate, median inter-spike interval, waveform asymmetry (peak asymmetry), and trough to after-hyperpolarization (AHP) duration. The peak asymmetry was defined as B-A/B+A where A is the first positive peak preceding the action potential trough and B is the first positive peak after the action potential trough (see inset in Fig. 2A) (2, 29). Trough-AHP duration was defined as the time from the action potential trough to B. These metrics were derived from data encompassing the entire recording session, excluding any post-clonidine injection intervals.

**Fig. 1.**
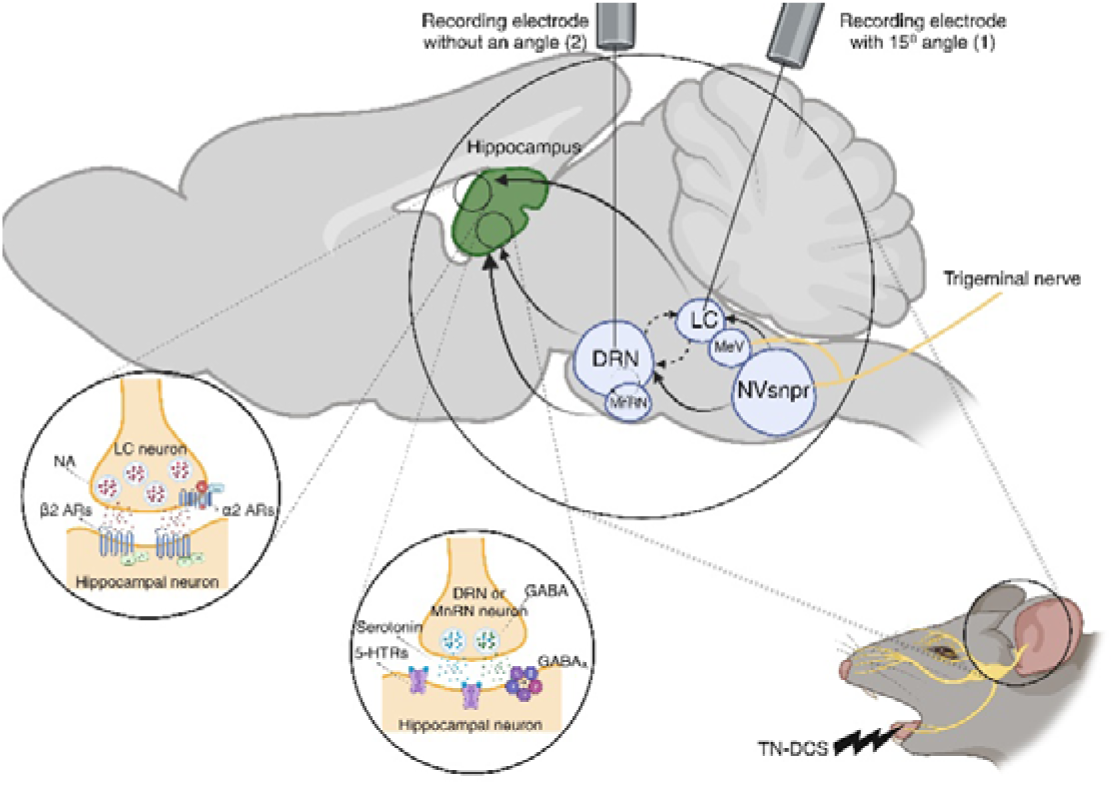
The schematic representation of the pathways implicated in mediating the effects of trigeminal nerve direct current stimulation (TN-DCS) on deep brainstem structures, subsequently influencing higher cortical areas like the hippocampus. We have previously shown that TN-DCS causes an increase in neuronal activity within the principal sensory (NVsnpr) and mesencephalic (MeV) nuclei of the trigeminal nerve (10). Here, we investigated if TN-DCS also affects activity in the locus coeruleus (LC), as well as the median and dorsal raphe nuclei (MnRN and DRN). This change in neural activity could potentially cause the release of serotonin and norepinephrine leading to alterations in higher cortical areas, such as the hippocampus (14). NA, noradrenaline; AR, adrenoreceptor; GABA, gamma-aminobutyric acid.

**Fig. 2.**
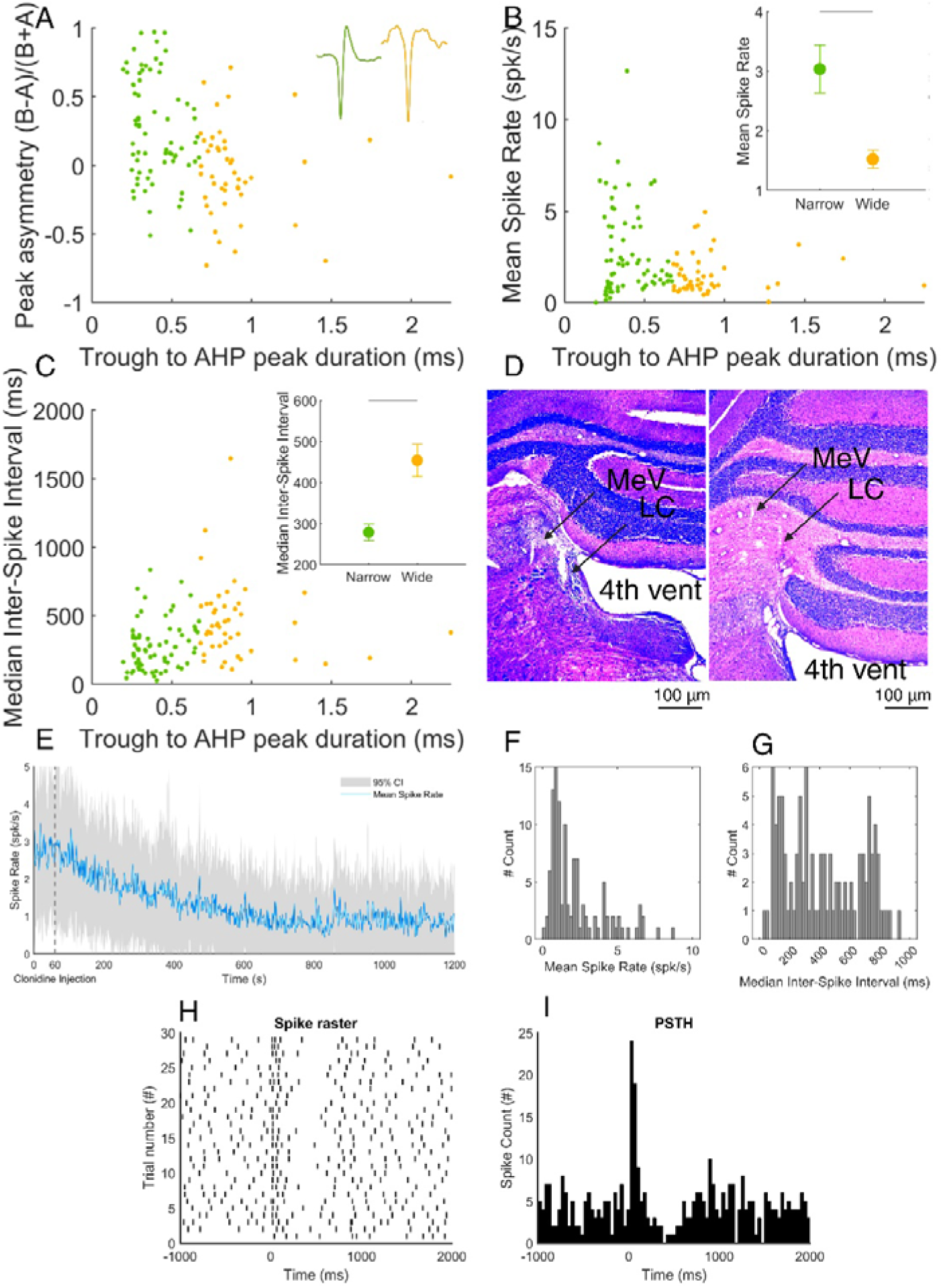
Classification of neurons in the LC based on electrophysiological characteristics (A, B, and C) and combined histological and electrophysiological verification of LC targeting (D, E, F, and G). Distinct populations of LC single units (green and yellow) were separable by waveform width and differed in spike rate and spatial distribution. (A) Units were separable based on the waveform duration and the amplitude of the after- hyperpolarization in relation to the first peak (n = 71 (narrow) and 46 (wide) units). The peak asymmetry was defined as B-A/B+A where A is the first positive peak preceding the action potential trough and B is the first positive peak after the action potential trough. (B) Scatterplots with the mean spike rate and (C) median inter- spike interval for each LC unit. The insets in (A), (B), and (C) show the average waveforms of two example units and the mean and standard deviation (SD) for each subset of units, respectively. The gray lines in the insets in (B) and (C) denote significant statistical differences between subsets of units. D) Nissl-stained coronal sections, each 40 μm thick, display the electrode trajectory made by the 15 μm probe angled at 15° within the coronal plane, marked by arrows. The slices are arranged from left (posterior) to right (anterior). Due to the 15° tilt, a single slice did not capture the entire electrode track. In every section, the top and bottom arrows indicate the electrode trajectories for the MeV and LC, respectively. E) Mean spike rate over time in the LC, initiated with a baseline recording for 60 seconds and administration of clonidine at a dosage of 0.05 mg/kg/i.p. afterward, and was recorded over 1200 seconds. The blue line represents the mean spike rate, while the grey-shaded areas indicate the %95 confidence intervals (CIs). F and G) histogram showing the mean spike rate (F) and median ISI for all neurons. H and I) Spike raster plot (H) and PSTH (I) showing a combined multi-unit spike train and spike count for each rat, respectively, created by merging the spike trains (F) and counts (G) of all recorded single units. These plots feature data obtained from 9 units in a rat. Zero in the X axis (Time) marks the moment the contralateral hind paw was stimulated at 3 mA. Following the stimulation (time 0) a collective brief burst of spikes can be seen in the population. This initial response is succeeded by an auto-inhibition phase mediated by noradrenaline autoinhibitory receptors on LC-NA neurons. PSTH, peristimulus time histogram; LC, locus coeruleus; MeV, the mesencephalic nucleus of the fifth cranial nerve; NA, noradrenergic.

In addition to electrophysiological criteria, the following anatomical landmarks helped locate the LC: 1) A region characterized by relative electrical silence just dorsomedial to the nucleus, aligning with the fourth ventricle; and 2) The MeV, situated just lateral to the LC, which exhibited cell activation in response to passive jaw stretch. To ensure the specificity of NA neuron spiking, clonidine (0.05 mg/kg i.p.; Sigma-Aldrich, Santa Cruz, USA) was administered at the end of the recording session (34, 35).

Here, the effects of TN-DCS on both the tonic and phasic activities of LC were evaluated. The baseline firing rate served as a proxy for the tonic activity, whereas the number of bursts, spikes per burst, burst duration, and inter-burst interval were metrics (named phasic activity metrics in the statistical model; see below) used to characterize the phasic activity. A burst was defined as two spikes that had an ISI of less than 0.08 seconds and ended with an ISI of more than 0.16 seconds, as previously mentioned (30).

### DRN and MnRN

To evaluate neuronal activity in the raphe nuclei, a 50 μm thick silicone probe with 32 channels covering a surface area of 1550 µm was used (Atlas Neuro, Leuven, Belgium, Model: E32+R-50-S1M-L20 NT). The location of the DRN and MnRN was determined using the Paxinos and Watson rat brain atlas as a reference. A burr hole craniotomy was performed using the same drill at coordinates AP: -7.92 mm, ML: 0.0 mm, and DV: (-6 to -7 mm for DRN and -7 to -9 mm for MnRN ventral to the skull surface). The following electroanatomical criteria were used to estimate the approximate position of the electrode tip: 1. An area characterized by a notable absence of electrical activity approximately 200 μm above the DRN, corresponding to the location of the Sylvian aqueduct, and 2. Neuronal activity in the target region identified by a broad action potential waveform and a slow, regular firing pattern (36).

### Stimulation and recording protocol

To activate the marginal branch of the TN in rats, a 1 cm² rectangular metal electrode was placed as an anode on the lower lip, while another electrode, serving as a cathode, was attached to the tail. Both electrodes were coated with Signa Gel (Parker Labs, New Jersey, USA) to ensure proper conductivity. DC stimulation was delivered according to the following protocol: 1 minute without stimulation (pre-stimulation), 1 minute of continuous DC stimulation (stimulation phase), and another 1 minute without stimulation (post- stimulation). This 3-minute stimulation protocol was then applied for each of the following amplitudes in the raphe nuclei (+0.25, +0.5, +1, +2, +3 mA) and the LC (+0.25, +0.5, +1, +2, +3, -0.25, -0.5, -1, -2, -3 mA). Each stimulation was continuously delivered once without repetition with an inter-stimulus washout period of 3 minutes.

The stimulation was administered using an AM 2200 analog current source (AM Systems, Sequim, WA), which received an analog voltage waveform from a data acquisition card (NI USB-6216, National Instruments, Austin, USA) operating at a sampling rate of 100 kHz and controlled by MATLAB 2014a (MathWorks, Natick, MA, USA). The signals from the 32 channels of the probe were amplified (x192), bandpass filtered (0.3 Hz–3 kHz), and digitized at 30 kHz and 16-bit resolution with a 32-channel recording head stage (RHD 32, Intan Technologies, Los Angeles, CA, USA). The probe data were continuously recorded on a PC via the Open Ephys GUI (https://open-ephys.org/).

To control for the specificity of TN activation, the ipsilateral TN was blocked at the end of several experiments (DRN, n=2; LC, n=8) by injecting 1-3 ml of 2% xylocaine (Xyl-M; VMD, Belgium) directly into the nerve and its facial branches. This procedure helped determine whether the responses observed in the brainstem nuclei were due to TN activation and not by volume conduction.

### Histology

After each experiment, rats underwent transcardial perfusion with phosphate-buffered saline (PBS) and fixation with a 4% paraformaldehyde (PFA) solution. Rat brains were then extracted, fixed in paraffin, sectioned, and subjected to cresyl violet (Nissl) staining to confirm the electrode placements.

### Data analysis and statistics

We used SpyKING CIRCUS 1.0.1 for spike sorting (37). The preprocessing steps included spike detection, spatial whitening, and principal component analysis (PCA). Accordingly, the data were clustered, merged, and template-matched. In the next step, the clustering was manually reviewed and modified using phy (https://github.com/cortex-lab/phy). For further analysis using MATLAB 2022a (MathWorks, Natick, MA, USA), unique single-unit clusters were chosen based on showing a V-shaped auto-correlogram (10).

For statistical analysis, tonic (i.e. mean spike rate) and phasic (i.e. number of bursts, inter-burst interval, number of spikes per burst, and burst duration in the LC; named as phasic activity metrics in the model, see below) activities were calculated using MATLAB. In line with our previous study and following the guidelines of Yu et al., 2022 (38), linear mixed-effect (LME) models were applied to evaluate the influence of stimulation amplitude and time epoch on spike rate for LC and RNs data (38). With raphe nuclei, rats, and neurons were regarded as random effects, whereas the stimulation amplitude, time epoch, and their interaction were handled as fixed effects. In the LC, rats, and neurons were regarded as random effects, whereas the stimulation amplitude, polarity, time epoch, and their interaction were handled as fixed effects in the following models:

1. Tonic activity:
2. For RN: Spike Rate ∼ Amplitude + Time epoch + Amplitude × Time epoch + (1|Neuron) + (1|Rat)
3. For LC: Spike Rate ∼ Amplitude + Time epoch + Polarity + Amplitude × Time epoch × Polarity + (1|Neuron) + (1|Rat)
4. Phasic activity:

For LC: phasic activity metrics ∼ Amplitude + Time epoch + Polarity + Amplitude × Time epoch × Polarity + (1|Neuron) + (1|Rat) 3) Xylocaine condition: Spike Rate ∼ Xylocaine × Time epoch + (1|Neuron) + (1|Rat).

The significance of the fixed factors and interactions in the model was evaluated using two-way ANOVA. A two-sided Wilcoxon signed rank test with a rigorous Bonferroni correction was performed to account for multiple comparisons during post-hoc analysis. This comprised three time epochs (pre, during, and post) and five stimulation amplitudes for the LC dataset, yielding fifteen multiple comparisons. For the DRN and MnRN datasets, we used three time epochs (pre, during, and post) and four stimulation amplitudes, resulting in twelve multiple comparisons for each nucleus. A significance threshold of 0.05 was taken into account in every comparison.

## Results

### Verification of LC targeting through histological and electrophysiological methods

Verification of LC targeting was achieved through several histological and electrophysiological methods (Fig. 2). Figure 2D displays two slides illustrating the trajectory of the electrode for both the MeV and LC. The MeV served as an electrophysiological approach to pinpoint the location of the LC. When the electrode was positioned within the MeV, mechanical stimulation of the rat’s whiskers led to a sharp increase in spike rate within the channels located inside the nucleus. Subsequently, the electrode was adjusted medially in steps of 50 to 100 μm until the electrophysiological signatures characteristic of the LC emerged. After the experiment, an injection of clonidine resulted in a marked reduction in the spike rate in the NA neurons of the LC, as depicted in Figure 2E. Fig. 2 F and G show the mean spike rate and median ISI for all neurons recorded from LC. Fig. 2H and I depict an example spike raster plot and peristimulus time histogram of 9 single units from one rat in response to foot shock, created by merging the spike trains of all recorded single units. After the stimulus (time 0), there was an initial rapid burst of spikes observed in the population. This immediate reaction was followed by an auto-inhibition phase facilitated by noradrenaline autoinhibitory receptors on LC-NA neurons.

### Effect of TN-DCS on tonic neural activity in the LC

From the LC, 57 neurons (n=7 rats) were isolated and studied for cathodic stimulation, and 117 neurons (n=11 rats) were isolated and studied for anodic stimulation. These results are shown in Fig. 3. We found a significant effect of TN-DCS amplitude (F(1 2205) = 14.499, p<0.001) on the spike rate in the LC. Additionally, there was a significant interaction between stimulation amplitude and time epoch (F(_2, 2205_) = 16.069, p = 1.1793e-07), stimulation amplitude and polarity (F(_2, 2205_) = 28.439, p = 1.0658e-07) as well as among stimulation amplitude, polarity, and time epoch (F_(2, 2205)_ = 9.028, p < 0.001). Conversely, the effect of the time epoch and polarity alone, the interaction between stimulation amplitude and polarity, and the interaction between time epoch and polarity did not reach statistical significance (p > 0.05 for all of these comparisons) (Fig. 3 A and B). Post-hoc testing showed that higher TN-DCS amplitudes caused higher spike rates in the LC during stimulation, with a significant p-value (p < 0.001) observed in comparisons between pre- and during-stimulation time epochs across +1, +2, +3, -1, -2 and -3 mA stimulation amplitudes. The increased stimulation amplitudes were also linked to higher Cohen’s d values, indicating a larger effect size. Please refer to Table 1 for detailed post-hoc test results.

**Fig. 3.**
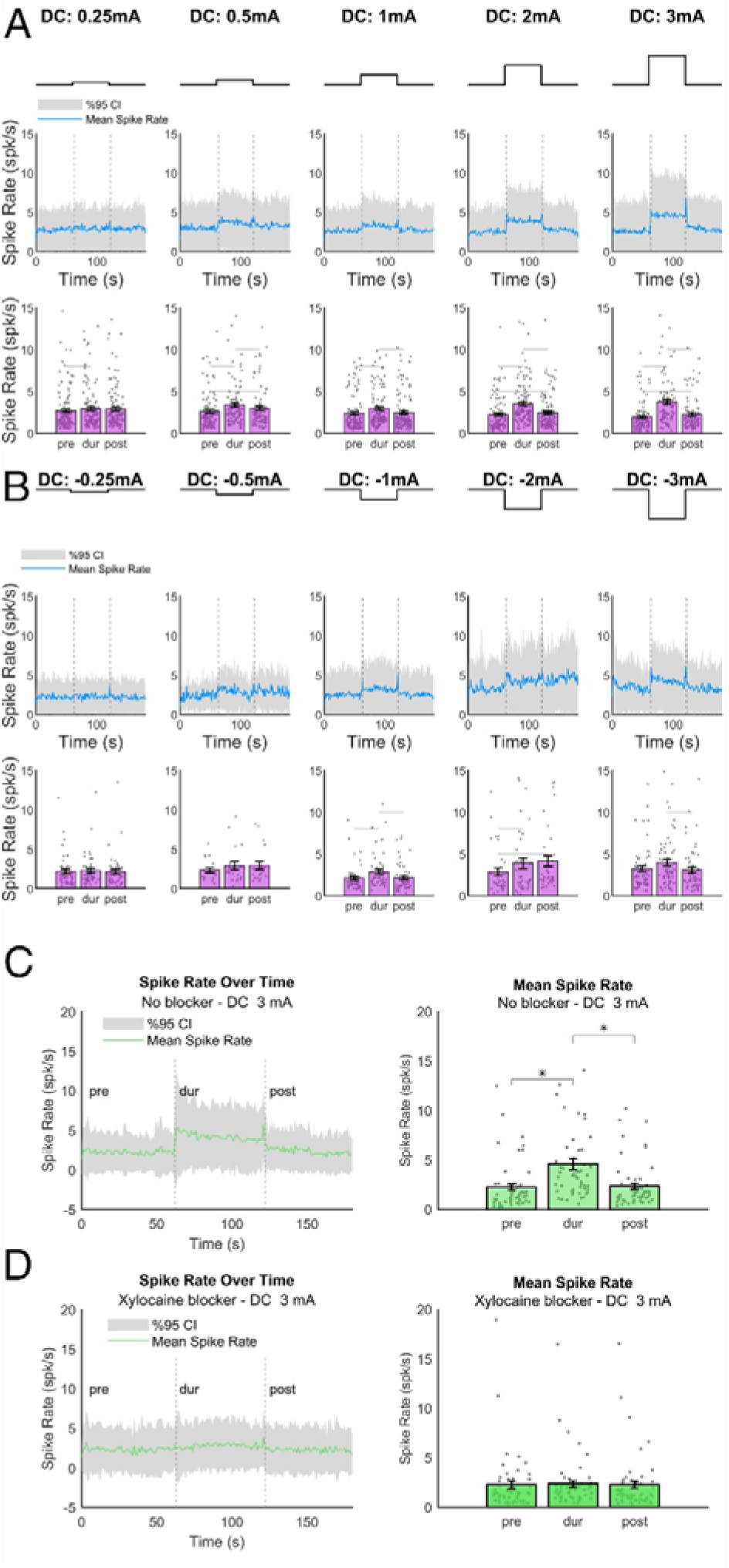
The effects of trigeminal nerve direct current stimulation (TN-DCS) and trigeminal nerve blockage via xylocaine on neuronal spike rates in rats’ locus coeruleus (LC). The study included 7 rats (57 neurons) for cathodic, 11 rats (117 neurons) for anodic stimulation, and 8 rats (51 neurons) for xylocaine conditions. A) Anodic TN-DCS: Significant increases in mean spike rate were observed at 1, 2, and 3 mA between pre- stimulation and during-stimulation time epochs, as well as between during-stimulation and post-stimulation time epochs. B) Cathodic TN-DCS: Significant increases in mean spike rate were observed at -1, -2, and -3 mA between pre-stimulation and during-stimulation time epochs, as well as between during-stimulation and post- stimulation time epochs. C and D) Xylocaine condition: During stimulation at maximum amplitude (3 mA) following xylocaine administration (D), there was no significant change in neuronal spike rate compared to the condition without xylocaine injection (C). The grey-shaded regions and blue lines in A and B, and green lines in C and D represent the 95% confidence intervals (CIs) and the mean spike rate over time for all neurons. Bar graphs with error bars in A, B, C, and D show the mean spike rate and standard deviation (SD) for the pre-stimulation, during-stimulation, and post-stimulation time epochs at various amplitudes. Each grey dot represents an individual neuron. Data were analyzed using a linear mixed-effects model, followed by a two-sided Wilcoxon signed-rank test with Bonferroni correction for multiple comparisons. Significant differences between time epochs are indicated by a grey horizontal line.

**Table 1.** An in-depth post-hoc evaluation designed to determine the impact of direct current trigeminal nerve stimulation (TN-DCS) on the spike rates of neurons in the locus coeruleus of rats (n=11). It includes detailed effect size data (Cohen’s d) along with corresponding *p*- and Z-values (mean deviations) across various anodic and cathodic amplitudes of TN-DCS. The analysis spans three different time epochs: pre- vs. during stimulation, post- vs. during stimulation, and pre- vs. post-stimulation in rats.

| Effect size measure | Anodic stimulation of LC |  |  |  |  |  |  |  |  |  |  |  |  |  |  |
| --- | --- | --- | --- | --- | --- | --- | --- | --- | --- | --- | --- | --- | --- | --- | --- |
|  | 0.25 mA |  |  | 0.5 mA |  |  | 1 mA |  |  | 2 mA |  |  | 3 mA |  |  |
|  | Pre-During | During-Post | Pre-Post | Pre-During | During-Post | Pre-Post | Pre-During | During-Post | Pre-Post | Pre-During | During-Post | Pre-Post | Pre-During | During-Post | Pre-Post |
| P value | 0.027 | 1 | 0.219 | 0.002 | 0.040 | 0.019 | 0.000 | 0.000 | 1 | 0.000 | 0.000 | 0.001 | 0.000 | 0.000 | 0.001 |
| Z value | -4.3652 | -1.564 | -2.359 | -5.337 | -4.212 | -4.444 | -6.662 | -5.820 | -1.460 | -8.405 | -8.202 | -3.901 | -7.242 | -6.699 | -3.810 |
| Cohen's d | 0.100 | 0.049 | 0.079 | 0.325 | 0.163 | 0.164 | 0.284 | 0.240 | 0.040 | 0.593 | 0.493 | 0.103 | 0.810 | 0.649 | 0.174 |
| Effect size measure | Cathodic stimulation of LC |  |  |  |  |  |  |  |  |  |  |  |  |  |  |
|  | -0.25 mA |  |  | -0.5 mA |  |  | -1 mA |  |  | -2 mA |  |  | -3 mA |  |  |
|  | Pre-During | During-Post | Pre-Post | Pre-During | During-Post | Pre-Post | Pre-During | During-Post | Pre-Post | Pre-During | During-Post | Pre-Post | Pre-During | During-Post | Pre-Post |
| P value | 1 | 1 | 1 | 0.375 | 1 | 0.118 | 0.092 | 0.017 | 1 | 0.047 | 1 | 0.022 | 1 | 0.01 | 1 |
| Z value | -1.639 | -0.939 | -0.395 | -2.153 | 0.165 | -2.579 | -2.663 | -4.735 | 0.773 | -2.880 | 0.823 | -4.683 | -1.721 | -3.925 | 0.642 |
| Cohen's d | 0.050 | 0.044 | 0.005 | 0.335 | 0.008 | 0.334 | 0.353 | 0.292 | 0.196 | 0.265 | -0.100 | 0.1374 | 0.268 | 0.344 | -0.090 |

### Effect of TN-DCS on tonic neural activity in the LC during TN xylocaine blockade

In our study, we isolated 51 neurons from the LC in 8 rats for the experiment with the xylocaine blocker. The group data from these neurons is shown in Fig 3C and Ds, indicating the lack of effect of TN-DCS when the xylocaine blocker was used. We found that the experimental time epoch (pre, during, or post), at the highest level of stimulation amplitude, tested i.e., 3 mA, did not significantly alter the spike rate following the administration of xylocaine (F_(2, 150)_ = 0.337, p = 0.713). This indicates that the previously observed effects of TN-DCS on the LC (see Fig. 3A), were caused by TN stimulation and not by stimulation volume conduction to the brainstem nuclei.

We also assessed the effects of TN-DCS on cells with various spike types in the LC (wide vs. narrow spike). However, no significant difference was found in the impact of TN-DCS on the spike rates of these two groups (Fig. 4).

**Fig. 4.**
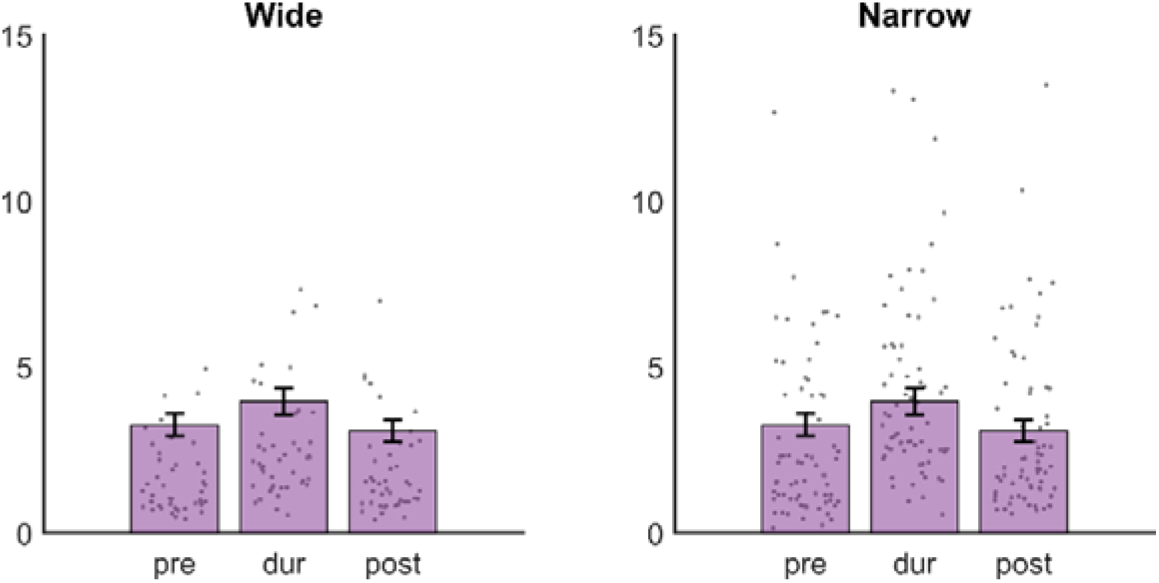
The effects of different amplitudes of trigeminal nerve direct current stimulation (TN-DCS) on neuronal spike rates in cells with various spike types (wide vs. narrow spike) in the locus coeruleus (LC) in rats (n = 70 (narrow) and 45 (wide) units). During anodic TN-DCS, no significant changes were observed in the mean spike rate at 3 mA between the pre- and during-time epochs, as well as between the during and post-time epochs. The bar graphs and error bars represent the mean spike rate and standard deviation (SD) for the pre-, during, and post-TN-DCS time epochs at 3 mA. Each grey dot signifies an individual neuron. The data underwent analysis using a linear mixed-effects model, followed by a two-sided Wilcoxon signed rank test with a rigorous Bonferroni correction for multiple comparisons.

### Effect of TN-DCS on phasic neuronal activity in LC

Four parameters were assessed to determine the effect of TN-DCS on phasic LC activity. As a result, it was discovered that amplitude, time epoch, the interaction between amplitude and time epoch, the interaction between amplitude and polarity, and the interaction between amplitude, time epoch, and polarity had a significant effect on the number of bursts in the LC (F_(1, 1854)_ = 9.026, *p* = 0.002, F_(2, 1854)_ = 5.676, *p* = 0.003, F_(2, 1854)_ = 16.608, *p* < 0.001, F_(1, 1854)_ = 16.537, *p* < 0.001, and F_(2, 1854)_ = 9.065, *p* < 0.001, respectively). However, for polarity, and the interaction between time epoch and polarity, this was not significant (*p* > 0.05 for all the comparisons) (Fig. 5A).

**Fig. 5.**
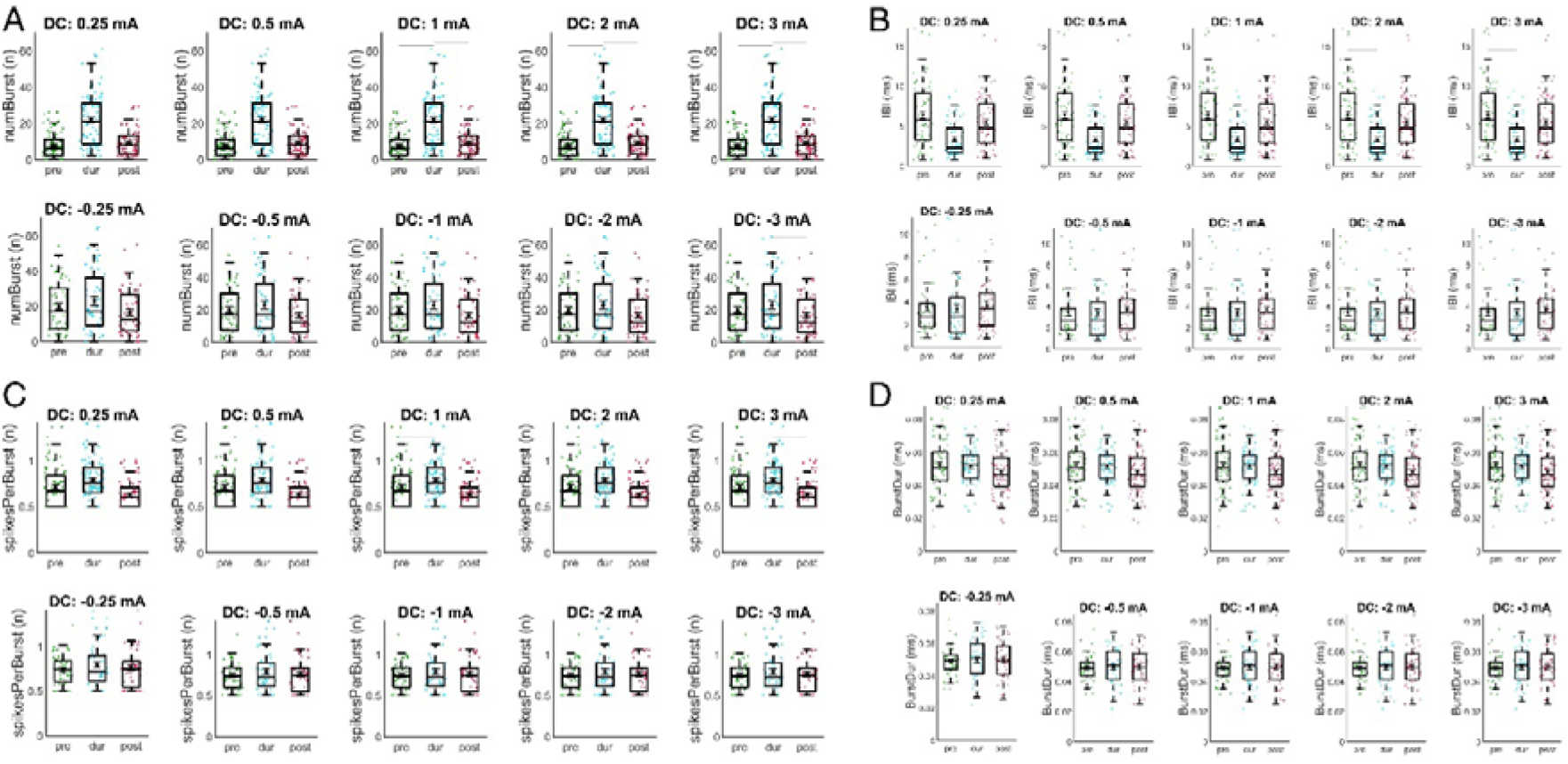
Effect of TN-DCS on phasic neuronal activity in the locus coeruleus (LC). This figure illustrates the effects of transcranial direct current stimulation (TN-DCS) on four parameters of phasic neuronal activity in the LC, analyzed across different current amplitudes and polarities. Data are presented for pre-stimulation (pre), during stimulation (dur), and post-stimulation (post) time epochs. Box plots showing (A) the number of bursts, (B) inter-burst interval (IBI), (C) the number of spikes per burst (spikesPerBurst), and (D) burst duration (BurstDur) before, during, and after TN-DCS at different current amplitudes (0.5 mA, 1 mA, 2 mA, 3 mA) for both positive (top row) and negative (bottom row) polarities. The difference seen in baselines of the number of bursts between anodic and cathodic conditions (A) could be due to the different number of animals used (7 rats cathodic vs. 11 rats for anodic stimulation. However, this difference between the baselines in anodic and cathodic polarities was not statistically significant (Mann-Whitney U test results: *p* = 0.054)). Each subplot shows median values and interquartile ranges, with individual data points indicated by colored dots, comparing the effects of TN-DCS across different amplitudes and polarities. Data were analyzed using a linear mixed- effects model, followed by a two-sided Wilcoxon signed-rank test with Bonferroni correction for multiple comparisons. Significant differences between time epochs are indicated by a grey horizontal line.

Furthermore, it was demonstrated that the inter-burst interval in the LC was significantly influenced by time epoch, the interaction between amplitude and time epoch, and the interaction between amplitude and polarity (F_(2, 1692)_ = 3.182, *p* = 0.041, F_(2, 1692)_ = 5.474, *p* = 0.004), and F_(1, 692)_ = 5.047, *p* = 0.024), respectively). The effects of amplitude, polarity, the interaction between polarity and time epoch, and the combination of polarity, amplitude, and time epoch, on the other hand, did not approach statistical significance (*p* > 0.05) (Fig. 5B).

The results showed that the following factors significantly affected the number of spikes per burst: time epoch (F_(2, 1803)_ = 3.336, p = 0.035) amplitude and polarity interaction (F_(1, 1803)_ = 4.862, p = 0.027), and amplitude and time epoch interaction (F_(2, 1803)_ = 3.158, p = 0.427). The effects of amplitude, polarity, their interaction, and the interaction between polarity, amplitude, and time epoch were not statistically significant, though (Fig. 5C).

Besides, it was found that polarity significantly affected the burst duration in the LC (F_(1, 1803)_ = 6.354, p = 0.011), but not time epoch or polarity. However, for the rest, this was not significant (p > 0.05 for all the comparisons) (Fig. 5D). Post hoc data are presented in Table 2.

**Table 2.**
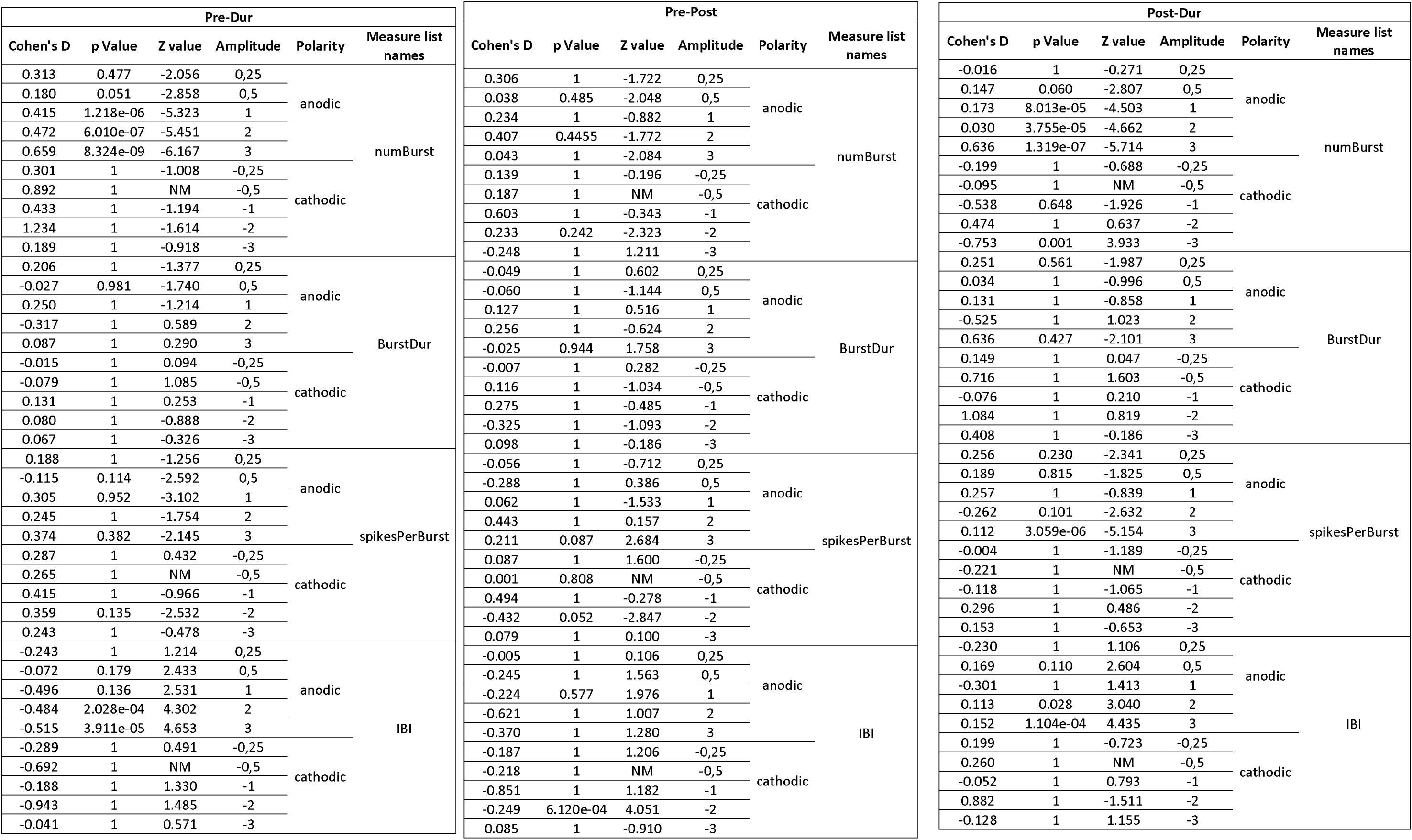
An in-depth post-hoc evaluation designed to determine the impact of direct current trigeminal nerve stimulation (TN-DCS) on the phasic activity of neurons in the locus coeruleus of rats (n=11). It includes detailed effect size data (Cohen’s d) along with corresponding p- and Z-values (mean deviations) across various anodic and cathodic amplitudes of TN-DCS. The analysis spans three different time epochs: pre- vs. during stimulation, post- vs. during stimulation, and pre- vs. post-stimulation in rats.

### Effect of TN-DCS on neuronal activity in DRN

205 single units were identified and isolated from the DRN. Fig. 6A collectively displays data for all DRN neurons, depicting their response to TN-DCS across different stimulation amplitude during the pre-, during-, and post-time epochs. The results of the post-hoc test are reported here. As polarity did not appear to have a major impact on the spike rate in LC, we only used anodic stimulation in RNs. We observed a significant effect of stimulation amplitude (F_(1, 2061)_ = 20.98, p = 0.0490), and the interaction between stimulation amplitude and time epoch (F_(2, 2061)_ = 5.016, p = 0.006) on spike rate. However, the effect of the time epoch on the spike rate was not statistically significant (F_(2, 2061)_ = 0.094, p = 0.909). The significant interaction between stimulation amplitude and time epoch suggests that higher stimulation amplitudes of TN-DCS caused increased spike rates in the DRN during stimulation as statistics showed a p < 0.001 for pre- vs. during-stimulation time epochs across all amplitudes. (Refer to Table 2 for comprehensive post-hoc testing details). Furthermore, higher stimulation amplitudes were associated with larger Cohen’s d values, indicating greater effect sizes (Table 3). Fig. 6 D and E represent the histological verification of DRN and MnRN targeting by the multichannel electrode. The DRN located just beneath the aqueduct (Aq) corresponds to the DRN, and further toward the distal part where the electrode tip is visible is the location of the MnRN.

**Fig. 6.**
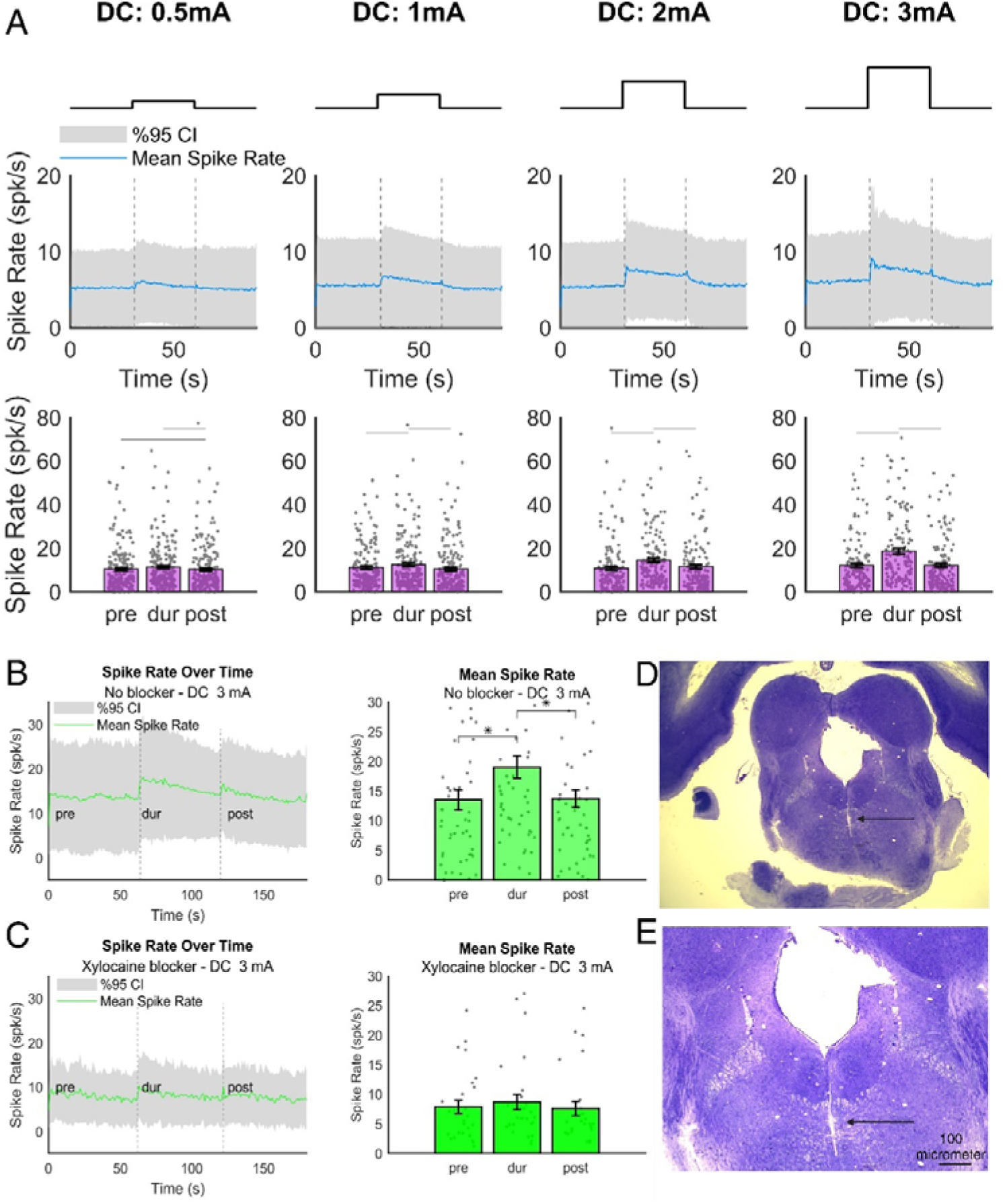
The effects of trigeminal nerve direct current stimulation (TN-DCS) with and without xylocaine on neuronal spike rates in the dorsal raphe nucleus (DRN) of rats. The experiment involved 9 rats (205 neurons) in the without-xylocaine condition and 2 rats (18 neurons) in the with-xylocaine condition. (A) Anodic TN-DCS significantly increased mean neuronal spike rates with higher stimulation amplitudes. (B, C) Post-xylocaine injection, 3 mA stimulation showed no significant difference in neuronal spike rates during the stimulation epoch compared to the pre-stimulation epoch. (D, E) Histological validation of electrode placement targeting the DRN and median raphe nucleus (MnRN) using a multichannel electrode at 1.25X (D) and 2.5X (E) magnifications. Black arrows indicate the electrode track along the midline, with the hole in the midline representing the Sylvian aqueduct. The y-axis represents spike rates in spk/s. Graphical representations include grey-shaded regions and blue lines in (A), and green lines in (B) and (C), showing 95% confidence intervals (CIs) and mean spike rates over time. Bar graphs with error bars in (A), (B), and (C) depict mean spike rates and standard deviations (SD) for pre-stimulation, during-stimulation, and post-stimulation epochs at various amplitudes. Each grey dot represents an individual neuron. Statistical analyses involved a linear mixed-effects model, followed by a two-sided Wilcoxon signed-rank test with Bonferroni correction for multiple comparisons. Significant differences between time epochs are indicated by a grey horizontal line.

**Table 3.**
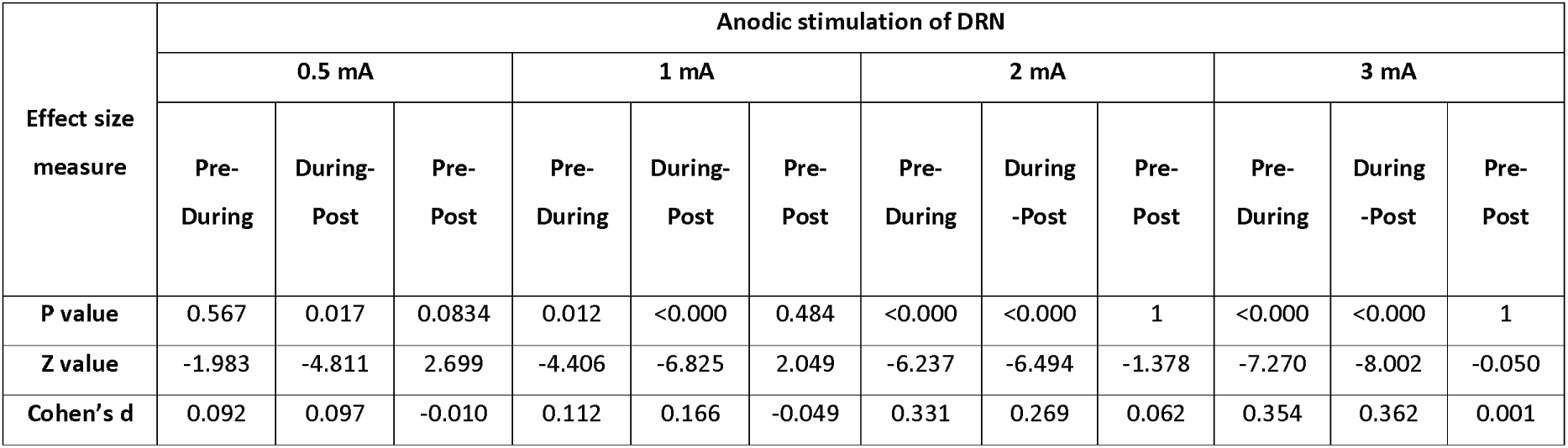
Post-hoc analyses to evaluate the magnitude of the impact of direct current trigeminal nerve stimulation (TN-DCS) on neuronal spike rates in the dorsal raphe nucleus in rats (n=9). This table presents effect size metrics (Cohen’s d), along with corresponding *p*- and Z-values (deviations from the mean) for varied anodic amplitudes of TN-DCS.

| Effect size measure | Anodic stimulation of DRN |  |  |  |  |  |  |  |  |  |  |  |
| --- | --- | --- | --- | --- | --- | --- | --- | --- | --- | --- | --- | --- |
|  | 0.5 mA |  |  | 1 mA |  |  | 2 mA |  |  | 3 mA |  |  |
|  | Pre-During | During-Post | Pre-Post | Pre-During | During-Post | Pre-Post | Pre-During | During-Post | Pre-Post | Pre-During | During-Post | Pre-Post |
| P value | 0.567 | 0.017 | 0.0834 | 0.012 | <0.000 | 0.484 | <0.000 | <0.000 | 1 | <0.000 | <0.000 | 1 |
| Z value | -1.983 | -4.811 | 2.699 | -4.406 | -6.825 | 2.049 | -6.237 | -6.494 | -1.378 | -7.270 | -8.002 | -0.050 |
| Cohen's d | 0.092 | 0.097 | -0.010 | 0.112 | 0.166 | -0.049 | 0.331 | 0.269 | 0.062 | 0.354 | 0.362 | 0.001 |

### Effect of TN-DCS on the spike rate in the DRN during TN xylocaine blockade

In the xylocaine blocker experiment, we isolated a group of 18 individual neurons from the DRN across 2 animals. Fig. 6B and C present the aggregated data for these neurons, depicting their responses to TN-DCS with the xylocaine blocker. Our findings indicated no significant effects for the experimental time epoch (pre, during, and post), showing no effect of the highest-tested stimulation amplitude on the spike rate after xylocaine injection (F_(2, 75)_ = 0.834, p = 0.438). This indicates that the effects observed during TN-DCS without xylocaine (see Fig 6A) were caused by TN stimulation, not by any stimulation volume conduction acting directly on the nuclei.

### Effect of TN-DCS on neuronal spike rate in MnRN

In the MnRN, 244 single units were successfully identified and isolated. Fig. 7A presents the data from all these neurons, showing their responses to TN-DCS over a range of stimulation amplitudes in pre-, during, and post-stimulation time epochs. Our analysis revealed no significant effect of stimulation amplitude (F(_1, 3660_) = 2.855, p = 0.091), time epoch (F(_2, 3660_) = 2.184, p = 0.112), nor an interaction between the two (F(_2, 3660_) = 0.354, p = 0.701) on the neuronal spike rate. This indicates that increasing the amplitude of TN-DCS did not cause a rise/decline in spike rates within the MnRN (p > 0.05 for comparisons between pre-stimulation and during-stimulation time epochs across all amplitudes).

**Fig. 7.**
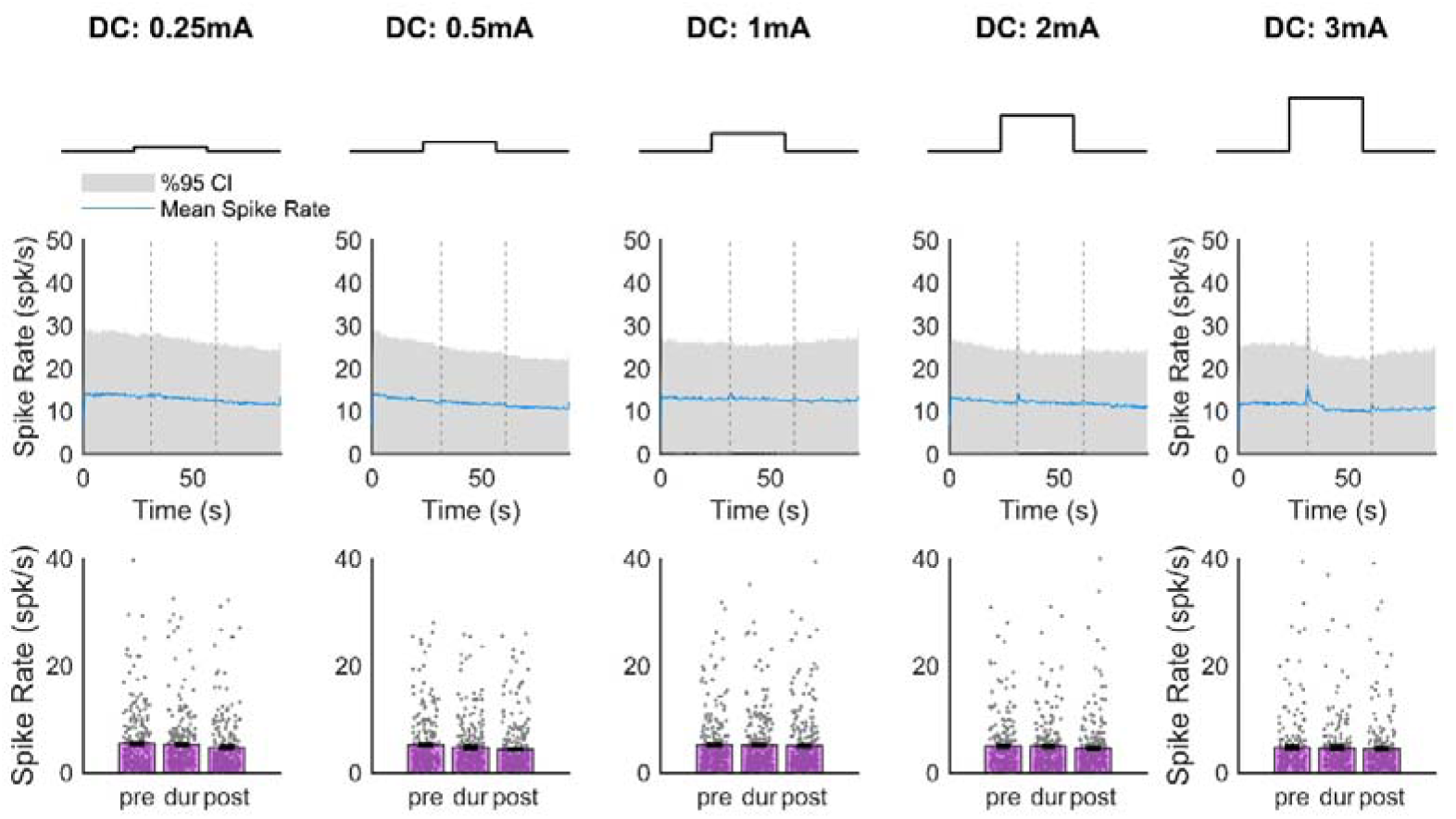
The effects of trigeminal nerve direct current stimulation (TN-DCS) with and without xylocaine on neuronal spike rates in the median raphe nucleus (MnRN) of rats. The experiment involved 5 rats (244 neurons). Anodic TN-DCS did not affect the mean neuronal spike rates at any tested amplitudes. The y-axis represents spike rates in spk/s. Graphical representations include grey-shaded regions and blue lines, showing 95% confidence intervals (CIs) and mean spike rates over time. Bar graphs with error bars depict mean spike rates and standard deviations (SD) for pre-stimulation, during-stimulation, and post-stimulation epochs at various amplitudes. Each grey dot represents an individual neuron. Statistical analyses involved a linear mixed-effects model, followed by a two-sided Wilcoxon signed-rank test with Bonferroni correction for multiple comparisons. Significant differences between time epochs are indicated by a grey horizontal line.

## Discussion

One of the principal challenges in the field of tDCS is the ambiguity surrounding its mechanism of action. While tDCS has been increasingly employed in clinical and experimental settings due to its non-invasive nature and ease of application, the underlying neurobiological processes that mediate its effects remain poorly understood. The prevailing theory suggests that tDCS modulates neuronal activity by subthreshold polarization of neuronal membrane potential, thereby influencing cortical excitability and plasticity (7, 39). However, the exact cellular and molecular pathways involved are not fully understood. This lack of clarity extends to its functional impacts, with studies showing variable effects on cognitive and behavioral outcomes (40). The heterogeneous responses observed across individuals and conditions further complicate the understanding of tDCS mechanisms (41). As such, a deeper investigation into the neurobiological mechanisms causing the effects of tDCS is crucial for optimizing its application and therapeutic efficacy.

Recent studies have introduced an intriguing perspective on the mechanism of tDCS, proposing that its effects might be at least partially mediated through transcutaneous mechanisms, i.e., the stimulation of trigeminal and occipital nerves (8). This hypothesis suggests that the traditionally assumed direct modulation of cortical neurons by tDCS might be complemented or overshadowed by co-stimulation of cranial/peripheral nerves in the scalp. Evidence supporting this theory stems from research indicating that scalp-applied electrical currents can activate superficial nerves, which could influence neural activity in the cortex (8, 42, 43). Our previous study that investigated the electrophysiological effects of TN-DCS on trigeminal nuclei showed that the first step in this pathway is viable, i.e. low amplitude direct currents activated the TN and increased neural spiking activity in both the NVsnpr and the MeV in rats. This effect was prevented by xylocaine injection to the ipsilateral nerve. Besides, tDCS without TN stimulation was not able to change neuronal activity in these nuclei implying that the increase in the mean firing rate and spike rate over time was due to the TN stimulation and not volume conduction via the skull or scalp (10). Previous neuroanatomical studies have provided evidence supporting the existence of reciprocal neural connections between the LC and the trigeminal sensory nuclei (e.g., NVsnpr) (44).

The results of the current study extend our earlier findings and elucidate a more extensive cranial nerve co-stimulation path. Following anodic and cathodic TN-DCS at various amplitudes (0.25-3 mA), a significant increase in tonic activity (mean spike rate and spike rate over time) was observed in the LC. In line with these results, a further investigation carried out in our laboratory investigated the hypothesis that TN-DCS can affect hippocampus function by means of a route involving the LC-NA system (14). The results of the study showed that TN-DCS at 1 mA markedly changed spike-field coherence and for up to 45 minutes, increased the hippocampus spike rate (14). In both of these studies, the blockage of the TN with xylocaine effectively prevented any increase in spike rate, indicating that these effects were not due to direct electric field stimulation in the brain. Moreover, when 1 mA TN-DCS was applied in conjunction with clonidine, which blocks the LC pathway, no increase in the hippocampal spike rate was observed. This suggests a crucial role of the LC-NA pathway in this process (14). These findings along with those of our previous studies signify the importance of transcutaneous mechanism via TN during tDCS (Fig. 1).

Additionally, our investigation demonstrated that TN-DCS significantly modulated the LC’s phasic activity (burst numbers, inter-burst intervals, and number of spikes per burst). Phasic, burst-like activity in LC neurons has been demonstrated to be traditionally linked to salient sensory events and behavioral approach changes (45). This implies that attention and arousal regulation is dependent on both the overall firing rate and the temporal structure of LC bursts. Linking prominent sensory inputs to appropriate behavioral responses may also depend on the exact timing and synchronization of phasic bursts throughout the LC population (45). If this exact burst timing is disturbed, attention and cognitive flexibility may be distorted. Furthermore, even while the LC system may be disrupted early in neurodegenerative illnesses, LC neurons’ basic burst-firing characteristics may be preserved as a compensatory mechanism, at least initially (46). In light of this, maintaining the exact temporal dynamics of LC burst firing during DC-TNS for conditions involving LC malfunction may represent a significant therapeutic target.

In direct relevance to these findings, studies conducted by Vanneste et al (8) on both healthy volunteers and rats revealed that applying a 1.5 mA direct current stimulation for 20 minutes to the greater occipital nerve (ON-tDCS) can improve memory. In humans, this memory enhancement was evaluated using indirect indicators of the LC-NA system’s activity, such as changes in pupil size, saliva α-amylase levels, and event-related brain potentials (ERPs). This effect was achieved by stimulating the ascending fibers of the ON, which connect to the LC and trigger the release of noradrenaline. The findings of this study highlighted the critical role of the LC-NA pathway in activating the hippocampus, changing the functional connections that contribute to the consolidation of memory (8).

In line with previous evidence on LC (2), we identified both narrow and wide spike types in our LC data set (see Fig. 2). Our analysis did not find any differences in the effects of TN-DCS on different spike types (see Fig. 4). However, more research is needed to specifically link narrow and wide spike types directly to potential different cell types (32, 47).

We hypothesize that the serotonergic pathway may also play an important role in mediating the effects of TN-DCS on higher cortical areas. Our study showed an increase in the mean spike rate and spike rate over time during acute TN-DCS at 1-3 mA in the DRN but not MnRN. The effects on the former were prevented by xylocaine injection to the ipsilateral nerve branches. The findings of a previous study suggested that a significant population of neurons within the DRN have connections with the trigeminal sensory complex and NVsnpr (48). Additionally, the results from another study revealed a structured, two-way connection between the DRN and LC, with notably robust projections extending from the DRN to the LC

(49). In line with that, Haddjeri et al., (50) found that the activity of 5-HT neurons in the DRN relies on continuous stimulation from noradrenergic input from the LC, which is facilitated through α_1_-adrenoceptors (50). Therefore, the increased activity of LC neurons we observed could represent one route, in addition to direct links from the trigeminal nuclei, contributing to the rise in spike rate within the DRN. Conversely, 5-HT neurons in the DRN exhibit regulatory effects on the functioning of LC-NA neurons primarily mediated by 5-HT_1A_ receptors (23, 50). It’s important to remember that the DRN consists of various neuron types, each with unique electrophysiological properties (24). Given these differences, these neurons might exhibit distinct responses when subjected separately to TN-DCS stimulation (50). Consequently, not all the spike rates we observed in the DRN, which are within the same ranges as those reported in the study by Zhou et al. (51), correspond with the much lower ranges typically associated with 5-HT neurons (52).

Our data showed that in response to TN_DCS, the DRN spike rate showed an initial increase followed by a steady drop, which was larger with higher amplitudes. The 5-HT_1A_ receptors, which serve an auto-receptor function on the soma of 5-HT neurons, are possibly the medium through which this is communicated. The control of 5-HT neurons’ firing activity mostly depends on the somatodendritic 5-HT_1A_ autoreceptors. Through a negative feedback mechanism, the firing activity of 5-HT neurons is reduced by an augmented activation of these autoreceptors caused by an increase in 5-HT (23). On the other hand, after prolonged stimulation, 5-HT_1A_ auto-receptors desensitize, and 5-HT neurons in the DRN resume their typical firing rate, as observed in the long-term VNS studies (53).

The available evidence indicates the existence of serotonergic connections originating from the DRN and projecting to the MnRN. However, the serotonergic inputs from the MnRN to the DRN predominantly terminate on DRN interneurons, likely of an inhibitory GABAergic nature (54, 55). Previous research has demonstrated that lesions in the MnRN increase theta rhythm in the hippocampus (54), a phenomenon linked to integrative processes crucial for higher cognitive functions (56). In our study, we observed no significant alteration (or a slight decrease, as depicted in Fig. 7 at higher amplitudes) in the mean spike rate and spike rate over time of MnRN neurons following acute TN-DCS at any of the tested amplitudes. This may be necessary for the effects observed in the study by Chen et al on the hippocampus where it was discovered that 1 mA TN-DCS significantly altered spike-field coherence and increased neuronal spike rate in the hippocampus throughout 45 minutes of recording (14).

Our study has several limitations. Firstly, we could not investigate the impact of TN-DCS on changes in neurotransmitters such as noradrenaline and serotonin within the target nuclei and associated cortical regions, including the hippocampus. The assessment of neurochemical alterations following TN-DCS could have provided complementary valuable insights (14). Secondly, the study lacked precise identification of the specific cell types recorded – this is an intrinsic limitation of extracellular electrophysiological recordings. Given the presence of various neuron types in LC, DRN, and MnRN, stimulating each of these neurons may yield different effects in higher cortical areas and consequently influence behavior. Thirdly, the intricate interaction between LC, DRN, and MnRN was not fully elucidated in this study, contributing to the complexity of understanding the exact dynamics during stimulation. Fourthly, our study could not investigate the effects of pulsed-TNS and its variations in frequencies, pulse widths, and amplitudes on the mentioned neural pathway. Fifthly, we collected the data while the animals were anesthetized. It is well known that the firing activity of assumed 5-HT neurons under anesthesia is similar to that of slow-wave sleep (about 1 Hz) (57). Therefore, recordings on awake rats would be required to show that the variance observed between the control and VNS-treated rats is not due to the anesthetic drug. Sixthly, our study assessed the effects of acute TN-DCS on neuronal activity in the DRN, MnRN, and LC. The impacts of chronic TN-DCS on these variables may be similar to or significantly different from those observed in acute settings. Finally, in our study, we did not include the analysis of serotonergic cell phasic activity in the DRN and MnRN due to challenges in isolating these cells based on established criteria for defining serotonergic neurons. The criteria typically used for isolating serotonergic cells include the presence of specific markers such as tryptophan hydroxylase (TPH) and the expression of serotonin transporter (SERT). Additionally, electrophysiological characteristics, such as firing patterns and response to pharmacological agents, are also considered (58, 59). However, we encountered difficulties in applying these criteria effectively, which limited our ability to accurately measure phasic activity that occurs specifically in these cells. By addressing these methodological challenges, future studies can gain deeper insights into the serotonergic modulation of brainstem pathways and their implications for tDCS efficacy.

In conclusion, the results of this study revealed an elevation in the tonic activity (i.e. mean spike rate and spike rate over time) within the DRN and LC at higher amplitudes (1-3 mA) following TN-DCS in healthy rats. Additionally, TN-DCS modulated phasic activity in the LC. Notably, these effects were abolished by the injection of xylocaine into the ipsilateral nerve branches that underwent stimulation. The increased phasic (LC) and tonic activities (LC and DRN) of neurons are of significance, considering these nuclei serve as pivotal centers for crucial neurotransmitters like norepinephrine and serotonin. These neurotransmitters, in turn, play a role in altering function in target cortical areas, such as the hippocampus and prefrontal cortex. Understanding these mechanisms sheds light on how TN-DCS and eventually tDCS may influence neuronal function in target regions and, subsequently, impact behavior. From a translational perspective, this study holds potential significance, as a profound comprehension of the mechanisms involved in the effects of TNS and tDCS can contribute to optimizing these widely-used therapeutic options in various pathophysiological conditions.

## Acknowledgments and funding

This study was supported by Fonds Wetenschappelijk Onderzoek [grant number G0B4520N] and the National Institutes of Health [grant number 1R01MH123508-01].

